# Agricultural pesticides do not suppress infection of *Biomphalaria* (Gastropoda) by *Schistosoma mansoni* (Trematoda)

**DOI:** 10.1101/2023.02.13.528426

**Authors:** Akbar A. Ganatra, Jeremias M. Becker, Naeem Shahid, Salim Kaneno, Henner Hollert, Matthias Liess, Eric L. Agola, Francis McOdimba, Ulrike Fillinger

## Abstract

**Background:** Schistosomiasis is a neglected tropical disease caused by trematodes of the genus *Schistosoma*. The pathogen is transmitted via freshwater snails. These snails indirectly benefit from agricultural pesticides which affect their enemy species. Pesticide exposure of surface waters may thus increase the risk of schistosomiasis transmission unless it also affects the pathogen.

**Methodology:** We tested the tolerance of the free-swimming infective life stages (miracidia and cercariae) of *Schistosoma mansoni* to the commonly applied insecticides diazinon and imidacloprid. Additionally, we investigated whether these pesticides decrease the ability of miracidia to infect and further develop as sporocysts within the host snail *Biomphalaria pfeifferi*.

**Principal findings:** Exposure to imidacloprid for 6 and 12 hours immobilized 50% of miracidia at 150 and 16 μg/L, respectively (nominal EC50); 50% of cercariae were immobilized at 403 and 284 μg/L. Diazinon immobilized 50% of miracidia at 51 and 21 μg/L after 6 and 12 hours; 50% of cercariae were immobilized at 25 and 13 μg/L. This insecticide tolerance is lower than those of the host snail *B. pfeifferi* but comparable to those of other commonly tested freshwater invertebrates. Exposure for up to 6 hours decreased the infectivity of miracidia at high sublethal concentrations (48.8 μg imidacloprid/L and 10.5 μg diazinon/L, i.e. 20 - 33 % of EC50) but not at lower concentrations commonly observed in the field (4.88 μg imidacloprid/L and 1.05 μg diazinon/L). The development of sporocysts within the snail host was not affected at any of these test concentrations.

**Conclusions:** Insecticides did not affect the performance of *S. mansoni* at environmentally relevant concentrations. Accordingly, pesticide exposure is likely to increase the risk of schistosomiasis transmission by increasing host snail abundance without affecting the pathogen. Our results illustrate how the ecological side effects of pesticides are linked to human health, emphasizing the need for appropriate mitigation measures.

**Author summary:** Schistosomiasis is a major public health problem in 51 countries worldwide. Transmission requires human contact with freshwater snails that act as intermediate hosts, releasing free-swimming life stages of the trematodes. The host snails are highly tolerant to agricultural pesticides used in plant protection products. Pesticides enter freshwaters via drift and runoff, and indirectly foster the spread of host snails via adverse effects on more sensitive competitor and predator species in the water. Increasing the abundance of intermediate hosts raises potential contact with the human definitive host while transmission of the pathogen is not affected.

Here we show that pesticides do not affect the ability of the trematode *Schistosoma mansoni* to infect and develop within its host snail *Biomphalaria pfeifferi* at environmentally relevant concentrations. Consequently, risk of schistosomiasis increases when pesticide pollution favours the proliferation of snail hosts whilst not negatively affecting the free-living parasites nor their development in their snail hosts. Measures to mitigate pesticide pollution of freshwaters should be a concern in public health programs to sustainably roll back schistosomiasis. Intersectional collaborations are required to bridge the gap between the agricultural and the public health sector in search of sustainable and safe methods of crop production.

## Introduction

Schistosomiasis remains a major public health problem in much of the world [1] despite the effort to eliminate this disease that is caused by the parasitic trematodes of the genus *Schistosoma* that use freshwater snails as intermediate hosts [2]. In western Kenya, which is considered a highly endemic area, two forms of the disease that are relevant to human health are present, intestinal schistosomiasis and urinary schistosomiasis [3]. Here, we focused on the intestinal schistosomes caused by *S. mansoni* which parasitize planorbid snails from the genus *Biomphalaria* [4]. The trematodes penetrate the snails as free-swimming larvae (miracidia) and undergo asexual reproduction as sporocysts within the snail before maturation into human-infecting free-swimming cercariae. This process takes about four weeks within the snail [2]. Agricultural activities have been shown to increase the risk of schistosomiasis by creating suitable habitats such as dams and irrigation canals which are suitable habitats for host snails whilst preventing their predators, such as river prawns, from accessing them [5]. Moreover, a recent study has shown that contamination of freshwater with agricultural pesticides can increase the likelihood of finding host snails in potential habitats, as well as the density of existing host snail populations [6]. Pesticides indirectly foster the highly tolerant host snails by affecting their more sensitive competitors [6] and predators [7,8]. In western Kenya, pesticide residues found in freshwater samples and within freshwater snails were most toxic to freshwater arthropods (*Daphnia magna*), followed by fish (*Onchorhynchus mykiss*) [9]. As several fish species are potential predators of the host snails [10,11,12,13], a reduction in their numbers would allow for increase in snail populations. The host snails are likely to benefit also from the observed toxicity to the macroinvertebrate community - represented by the test species *Daphnia magna* - that include both potential predators and competitors of the snails [7,6,14]. Consequently, pesticide pollution may alter the risk of schistosomiasis transmission by supporting increased numbers of intermediate host snails. However, assessing effects of pesticide pollution on the risk of schistosomiasis requires also understanding how pesticides might affect the free-swimming life-stages of the pathogen *Schistosoma* itself.

In this study, we investigated effects of the insecticides imidacloprid and diazinon on larval stages of *S. mansoni* and on their interaction with the intermediate host snail *Biomphalaria pfeifferi*. Both insecticides are common in freshwater bodies in western Kenya [9]. Both insecticides were also shown to be potentially beneficial to host snails by being more lethal to every other macroinvertebrate species that has been collected from western Kenyan freshwater bodies and tested [6]. Miracidia of *S. mansoni* hatch from eggs excreted with human faeces and need to find a suitable snail host within 24 hours. During this period, the miracidia have limited energy reserves but they can chemotactically navigate in water to find their host [15,16]. We assessed whether miracidia host seeking is affected at high sublethal concentrations serving as positive control (20 - 33 % of the identified acute EC50, i.e. the median effective concentration that immobilizes half of the test organisms) and at environmentally relevant concentrations (2 – 3 % of the EC50). The lower test concentrations represent the upper range of pesticide toxicity that has been observed in rural freshwater bodies in western Kenya (Kandie *et al.*. 2020) and also around the world (i.e. a toxic unit, defined as 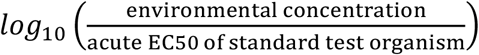, of 0 to −1 [6,14,17]. After successful infection, a single miracidium produces thousands of sporocysts within a snail that are then shed as cercariae and infect humans via dermal contact. Therefore, maturation and replication of *Schistosoma* relies on nutrient supply from the snail [18]. As a consequence, sporocysts are indirectly susceptible to environmental conditions such as pesticides that affect the energetic reserves but also the immune system of the snails [19]. Additionally, sporocysts may be directly affected by pesticide residues that enter the body of snails. As such, we assessed whether the maturation of miracidia to cercariae within the host snail *Biomphalaria pfeifferi* is disrupted when the host snail is exposed to pesticide pollution after being parasitized.

## Materials and methods

### Study location

All experiments were conducted at the International Centre for Insect Physiology and Ecology (*icipe*) Thomas Odhiambo Campus (TOC), Mbita, western Kenya.

### Snail collection and rearing

*Biomphalaria* snails were collected from the shores of Lake Victoria with a snail catcher and a pool net. Species were identified with a field identification key [20]. Collected snails were placed in open plastic containers along with some vegetation from the collection site for shade and cooling during transport. No water was provided during transport to avoid excess mortality due to warming. In the laboratory, snails were placed in large plastic tubs (45 x 35 x 28 cm) with 5 litres of lake water and reared with boiled kale (*Brassica oleracea L*) and tropical fish food. These tubs were kept at ambient conditions in a greenhouse with netting screened walls at *icipe* TOC. The day after collection, snails were screened for infection with a *Schistosoma* parasite by placing them individually in 24 well plates and exposing them to indirect sunlight for two hours to cause shedding of cercariae., similar to what was done by Opisa *et al*. [21]. After two hours, the well plates were observed under a dissecting microscope (Zeiss AxioCam5 100–400x) for *Schistosoma* cercariae which would indicate which snails were infected. Infected snails were separated and subsequently used to produce cercariae for experiments, but were otherwise reared similarly to uninfected snails in aerated, dechlorinated water and fed with boiled kales. Uninfected snails were reared for an additional 5 weeks before rechecking for cercarial shedding, after which uninfected snails were considered fit for experiments that required infection such as the miracidia host seeking and sporocyst development assays.

### *Schistosoma* cercariae collection

Cercariae were obtained from *Schistosoma* positive snails. On the days of experiments, the snails were placed in 24 well plates under artificial light at 9 am to allow for cercariae shedding for two hours before setting up assays. The well plates were then observed under a microscope (Zeiss AxioCam5 100–400x), the snail was removed, and cercariae were pipetted into the test containers for experiments as described in the acute toxicity tests section below. After the experiment, the snails used to shed the cercariae were placed in a freezer to kill them.

### *Schistosoma* miracidia collection

Miracidia were obtained from *Schistosoma* eggs extracted from stool samples obtained from primary school children with due consent from both parent and child, and ethical approval from the relevant national authorizing body. Over the course of the experiment, 145 children were recruited for screening from Kombe (–0.440028, 34.220040) and Wasulwa A (-0.435084, 34.211620) villages in Homa Bay County and Katito (–0.314557, 35.006869), Kisumu County. Stool samples of standard size - 41 mg, or described as about the size of a pea, were obtained from the children and tested for infection through the Kato-Katz method [1, 22]. Briefly, the stool was placed on a template that approximates the sample to about 43 mg per slide. The samples were then pressed with a cellophane strip coated with Malachite green dye, and slides were observed under a compound microscope (Axiocam ERc5s at 400x magnification) for *S. mansoni* eggs. The number of eggs per gram (epg) for each slide was counted. Forty-five children were found to be positive, of which those with 200 epg or more were recruited to provide additional stool samples to supply eggs as a source of eggs for miracidia while those with low egg burdens were immediately treated with praziquantel, according to the Kenya Government Ministry of Health guidelines using a Ugandan-model dose pole [23]. The children who provided samples were treated afterwards with a single dose of praziquantel (40mg/kg). All treatments were done under the supervision of a qualified and competent clinician. The stool was collected in plastic containers with lids sealed with cling film. *Schistosoma* eggs were isolated from the stool sample by passing it through a series of sieves of different pore sizes (212, 180, 150, 45μms) using 8.5% saline solution to harvest eggs and ensure they do not hatch. The isolated eggs were stored overnight in falcon tubes with saline, and experiments were conducted the day after egg collection to ensure their viability was not affected by storage time or overexposure to cold temperatures. When miracidia were needed for experiments, the falcon tubes with the eggs were poured into 5 litres of bottled water in a large conical flask. The flask was left on a bench near a window for 2 hours to allow the ova to hatch into miracidia. Afterwards, the flask was covered with a piece of aluminium foil so that the phototropic miracidia swam up to the water surface, where they were collected and transferred into a petri dish and utilised in the experiment.

### Insecticides

We tested the effects of two insecticides with different modes of action, the neonicotinoid imidacloprid and the organophosphate diazinon. Both compounds are among those that typically drive the overall risk of agricultural pesticides to freshwater invertebrates in the study area [9]. Imidacloprid was provided with the formulated plant protection product Loyalty® 700 WDG (distributed by Greenlife Crop Protection Africa, Nairobi; manufactured by Shandong United Pesticide Industry China) containing 700g imidacloprid per kg as the active ingredient. Diazinon was provided as the product Diazol® 60 EC (emulsified concentration, repacked and distributed by Laibuta Chemicals Ltd, Nairobi, an insecticide of the chloronicotinyl class; and containing an insecticide that contains diazinon 600g per kg, an organophosphate, as an active ingredient. Both formulated products are commonly sold in the study area. Fresh stock solutions based on the required active ingredient concentration were prepared the night before the experiments. The stock solutions were then left to stir overnight in amber glass bottles covered with foil and used to produce the remaining concentrations through dilutions on the morning of experimentation.

### Miracidia and cercariae acute toxicity assay

Immobilization of the miracidia and cercariae was recorded after constant exposure to imidacloprid and diazinon for 24 hours under a dissecting microscope (Zeiss AxioCam5 100-400x). Observations were made beginning at one and a half, three, six, 12 and finally 24 hours after exposure. As the test organisms had a mortality of 100% at 24 hours as miracidia, later analysis was limited to 12 hours. Test concentrations were based on preliminary experiments such that they covered the range of 5-95% mortality to estimate the median effective concentration required to immobilize 50% of the individuals (EC50) in 12 hours. Miracidia were tested at ambient temperate conditions (approx. 25 °C) with the following nominal test concentrations of both imidacloprid and diazinon: control (no compounds), 1, 4, 14, 55 and 209 μg/L). Each petri dish contained ten miracidia in 2 ml test concentration. Cercariae were tested with the following concentrations of both pesticides: control (no compounds), 1, 4, 14, 55, 209 and 792 μg/L. Each petri dish contained ten cercariae in 2 ml test concentration. Cercariae in the first petri dishes were tested at approx. 25°C like the miracidia. However, we observed high mortality in the controls so that subsequent tests with cercariae were done in a temperature-controlled room at 18°C. Tests were done on triplicate on three separate days, except for cercariae exposed to imidacloprid, to which a fourth day with three replicates was also done (Tab. S1 in supplementary). To account for potential host-mediated variability, the tested cercariae were collected and mixed from different snails and miracidia were collected and mixed from egg batches from different children.

### Miracidia host-seeking assay

We exposed miracidia to different concentrations of pesticides for either two, four, or six hours before allowing them access to a snail host. Using a pipette under a dissecting microscope, we distributed 1080 miracidia to six petri dishes (60 x 20 mm, PYREX 1480102D). Two petri dishes served as control, while each of the other dishes contained one of the following nominal test concentrations: 4.88 μg/L imidacloprid (~3% of the 6h EC50 for miracidia), 48.8 μg/L imidacloprid (~33% of the 6h EC50), 1.05 μg/L diazinon (~2% of the 6h EC50 for miracidia), and 10.5 μg/L diazinon (20% of the 6h EC50) such that each concentration was occupied by 180 miracidia. After pesticide exposure for two, four and six hours, respectively, 60 miracidia were collected from each petri dish and distributed into twelve 100 ml borosilicate crystallizing glasses (70 x 40mm, PYREX, Fig. 1). Each cup contained 70 ml bottled water and a single non-contaminated *Biomphalaria pfeifferi* snail, such that each of the twelve snails per pesticide, concentration and exposure time was exposed to five miracidia. The miracidia were given six hours to infect their host snail. The snails were then removed from the oviposition cup and washed with bottled water. They were reared in round plastic tubs (48cm diameter) with 2.5 L of lake water in the screenhouse as above, for three days to ensure successful infection has taken hold within the snail. After the three days, the snails were frozen and stored for molecular analysis to confirm successful penetration by the miracidia as described below.

**Fig 1:**
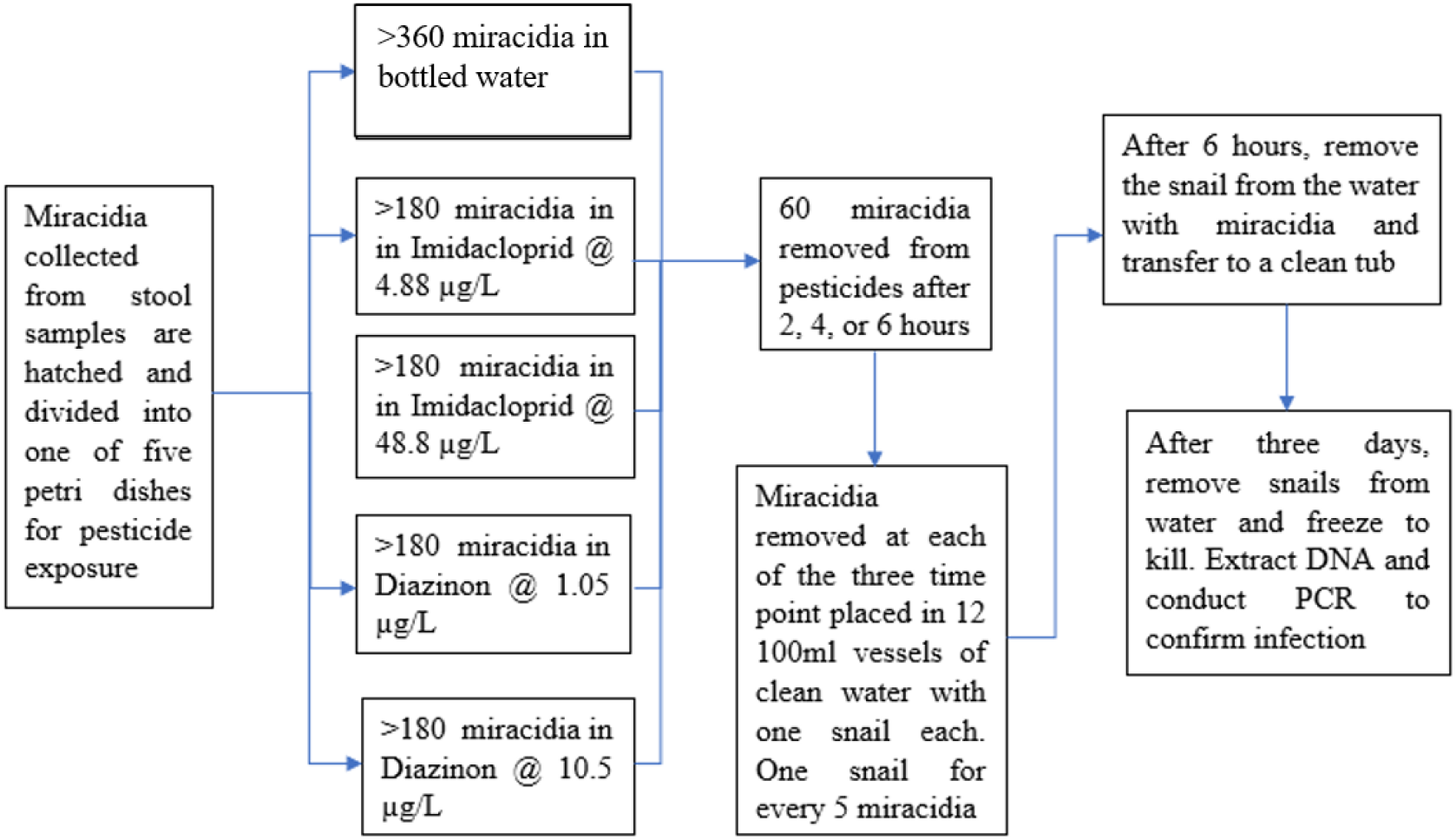
Flowchart for the miracidia host seeking assay.

### Sporocyst development assay

To test the effect of pesticide exposure on the growth and maturation of sporocysts, we first exposed 240 of the non-infected snails with fresh miracidia, with the aim to infect them. This was done by placing each snail with five miracidia in a glass dish with 100 ml of bottled water for six hours. We used bottled water to provide optimum conditions for the miracidia to penetrate and infect the snails. After infection, the snails were distributed into six 48 cm circular plastic tubs containing 40 snails each. The tubs were then randomly assigned to pesticide test concentrations similar to those in the miracidia host-seeking assay (as above): two controls, 4.88 μg/L imidacloprid (~3% the 6h EC50 for miracidia and 48.8 μg/L imidacloprid (33% the average 6h EC50 for miracidia), 1.05 μg/L diazinon (~2% the 6h EC50) and 10.5 μg/L diazinon (20% EC50). All test concentrations were below 0.01% of the 24h acute median lethal concentration (LC50) for *B. pfeifferi* (Becker *et al*., 2020). To mimic pulse exposure in the field, the infected snails were exposed to the described nominal pesticide concentrations at ambient temperature in glass bowls (48 cm diameter) for 24 hours once a week, beginning three days post infection (Fig 2). After pesticide exposure the snails were washed with lake water before being placed in their original tubs with lake water. The snails were otherwise reared in lake water, as above, that was changed weekly and fed boiled kale. Two of the tubs served as controls, the remaining tubs were randomly assigned to one of the following pesticide concentrations: 4.88 μg/L imidacloprid, 48.8 μg/L imidacloprid, 1.05 μg/L diazinon, and 10.5 μg/L diazinon. All test concentrations were below 0.01% of the 24h acute median lethal concentration (LC50) for *B. pfeifferi* [6]. Test concentrations resembled 2-3% and 20-33% of the average 6h EC50 of imidacloprid and diazinon for miracidia. Once a week, beginning three days post-infection, the snails were removed from their tubs of lake water and exposed to the described pesticide concentrations for 24 hours in glass bowls (48 cm diameter). After 24 hours, the snails were washed with lake water before being placed back in their original tubs with lake water. Beginning four weeks after the first exposure, the snails were checked every two days for cercariae shedding by exposing them to artificial light and observing them under a compound microscope. All positive snails were immediately removed for storage in 70% ethanol. Dead snails were also collected in 70% ethanol. Molecular screening was done on all stored samples for *Schistosoma* DNA to confirm the infection status as well as to detect prepatent infections that did not lead to cercarial shedding. The experiment concluded when all snails had died or been collected.

**Fig 2:**
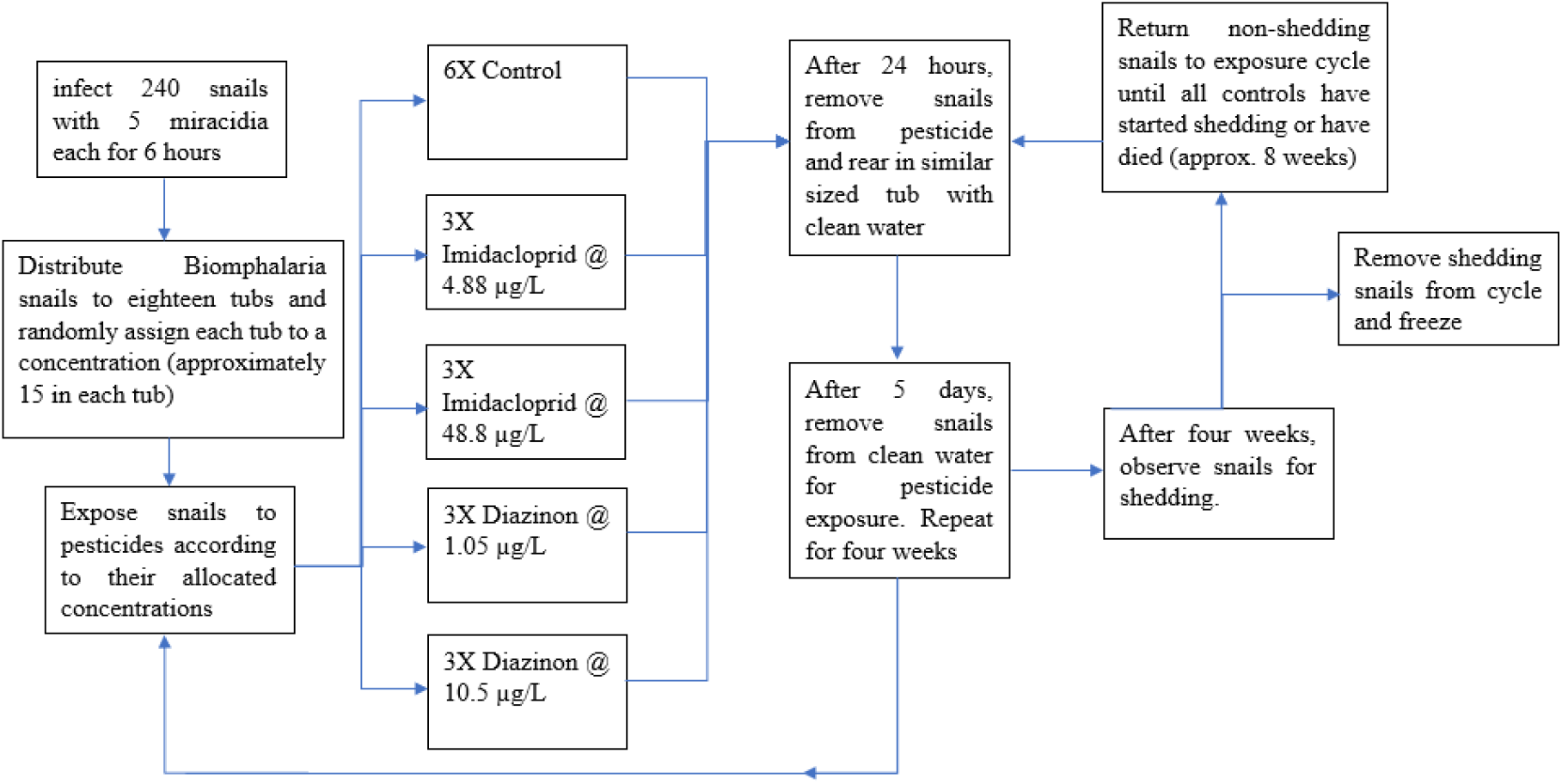
Flowchart of the sporocyst development assay.

### Molecular analysis

To ensure penetration of snails by miracidia had occurred, snails were tested for *Schistosoma* infection using polymerase chain reaction (PCR) assays to amplify any *Schistosoma* DNA within the snail as done in Sady *et al*., [. At point of testing, snails were removed from the freezer and the soft body extracted from the shell using forceps, and cut into small pieces. The bodies were then transferred to an Eppendorf tube and homogenised using a motorised homogeniser, and DNA was extracted using standard protocols [24]. The DNA obtained was then amplified using conventional PCR [25] using the following primers: ShbmF (5’- TTTTTTGGTCATCCTGAGGTGTAT-3’), ShR (5’- TGATAATCAATGACCCTGCAATAA-3’) and SmR 5’- TGCAGATAAAGCCACCCCTGTG-3’). Briefly, a total volume of 50 μl containing 10 mM Tris-HCl (pH 9.0), 50 mM KCl, 0.1% Triton® X-100, 200 μM dNTP (Promega, Madison, WI, USA), 2.5mM MgCl_2_, 0.2 μM of each primer, 1 unit of *Taq* polymerase (Promega, Madison, WI, USA), and approximately 75 ng of schistosome genomic DNA. The thermal cycling profile included an initial denaturation step at 94 °C for 5 min, followed by 35 cycles of 45 s at 94 °C, 45 s at 58 °C, 30 s at 72 °C and a final step of 7 min at 72 °C using a GeneAmp 2400 (Applied Biosystems, Foster City, CA, USA) thermal cycler. Amplicons were electrophoresed in a 10% agarose gel, stained with ethidium bromide and visualised in a UV chamber (ngenius syngene bio imaging. A 100bp DNA ladder was used to determine the product sizes. Presence of *Schistosoma* DNA was confirmed by a band on the agarose gel at 250 bps, indicating miracidia penetration of a snail (Fig S1)

### Data analysis

Datasets were analysed using RStudio for Windows (version 4.1,1, 2021-08-10) and R for Windows (software R 3.6.2)

R Core Team 2020 [26]. When analysing the acute toxicity tests with miracidia and cercariae, only replicate tests in which >60% of organisms in the controls survived were considered. Moreover, when two out of three replicates from the same day showed high mortality in the controls, all three replicates were discarded. Ultimately, this resulted in two to five replicate petri dishes per concentration being analysed at 12 hour (diazinon miracidia:cercariae = 5:5; imidacloprid miracidia:cercariae = 2:5), and six to ten replicate petri dishes per concentration analysed for six hours (diazinon miracidia:cercariae = 8:6; imidacloprid miracidia:cercariae = 9:10)(Supplementary Tab. S1 for details on which replicates were used).

First, we tested whether the different test temperatures had a significant effect on the observed concentration vs. immobilization relationship for cercariae. Therefore, we fitted a quasi-binomial generalized linear model (GLM) to the data from tests with both pesticides imidacloprid and diazinon together. We specified the following effects (explanatory variables) incl. all their interaction terms: pesticide identity, pesticide concentration, exposure time (6 and 12 h) and temperature. To improve model diagnostics, pesticide concentration was log-transformed and 0.5 (half of the lowest test concentration) was added to avoid negative infinite values for the control. We used a probit link function which provided the best fit. Then we applied backward selection by successively removing all effect terms that were considered non-significant (*p* > 0.05) based on χ^2^-tests for the increase in residual deviance due to effect term removal [27]. The resulting minimal adequate model contained only pesticide identity, pesticide concentration, exposure time, and the two-way interaction of pesticide identity and pesticide concentration, but not temperature (see results). Therefore, temperature was not considered relevant in the following analysis.

We estimated the median effective concentration which increased immobilization of miracidia and cercariae by 50% (EC50) from non-linear regression using the drc package 3.0-1. While it is more difficult with these models to test complex effect interactions, model fitting is more flexible as the upper limit of the fitted dose-response curve is variable and can be estimated from survival in the controls. This way, the EC50 may be estimated with higher precision as compared to GLMs. We used separate three-parameter binomial log-logistic models for each pesticide and life stage (miracidia and cercariae). Only in case a three-parameter model provided a bad fit (assessed visually), we used a five-parameter model instead; this was the case for the analyses of the effects of diazinon on miracidia after 6 h and on cercariae after 12 h. The five-parameter models included an additional parameter for the lower boundary of the fitted log(dose)-response curve (which was pre-set to zero) and a shape parameter to enable the fitting of asymmetric log(dose)-response curves. Because the different test temperatures for cercariae showed no significant effect on the log(dose)-response relationship in the GLM (see above), we did not differentiate according to temperature in the non-linear models.

The infection success of miracidia on snails was analysed using a binomial generalized linear model with a probit link function. The model contained pesticide identity, pesticide treatment (control, low and high concentration) and exposure time. Pesticide treatment was specified as a categorical factor and not as a numeric variable to test for significant effects at the limited number of test concentrations rather than fitting a full dose response curve. The model was reduced using backward selection as described above; only the main effect of pesticide treatment remained in the final model. The model was analysed using the R packages MASS 7.3-51.5 and effects 4.1-4. The different pesticide treatment levels in the model were compared using likelihood ratio χ2 tests with the phia package 0.2-1.

Pesticide effects on the sporocyst development were analysed with a similar generalized linear modelling approach. The initial model contained pesticide identity, pesticide treatment and their interaction as fixed effects and a logit link function. After backward selection, no effect terms remained in the minimum adequate model (see results). Pesticide effects on the sporocyst development and on the mortality of snails were analysed with a similar generalized linear modelling approach. The initial models contained pesticide identity, pesticide treatment and their interaction as fixed effects and a logit link function. After backward selection, no effect terms remained in the minimum adequate models for sporocyst development and for snail mortality (see results).

### Ethical clearance

Ethical Clearance was granted from the Kenya Medical Research Institute’s (KEMRI) Scientific and Ethical Review Unit (SERU) to collect stool samples from schoolchildren to obtain miracidia for experiments (KEMRI/SERU/CBRD/194/3836).

## Results

### Miracidia and cercariae acute toxicity tests

The life span of miracidia and cercariae is limited to approximately 24 hours, such that we observed a drastic reduction in control survival of miracidia after 12 hours. Thus, we limited the analyses of EC50s to effects after exposure for 6 and 12 hours rather than for 24 hours which is more common in ecotoxicological testing of other species. Based on non-linear regression analysis,the exposure to imidacloprid immobilized 50% of miracidia at 149.7 (12.8 – 1746) μg/L after 6 hours (mean ± 95% confidence interval) and at 15.6 (7.8 – 31.3) μg/L after 12 hours (Fig 3). Exposure to diazinon immobilized 50% of miracidia at 51.3 (37.9 – 69.4) μg/L after 6 hours and at 20.9 (14.7 – 29.8) μg/L after 12 hours.

**Fig 3.**
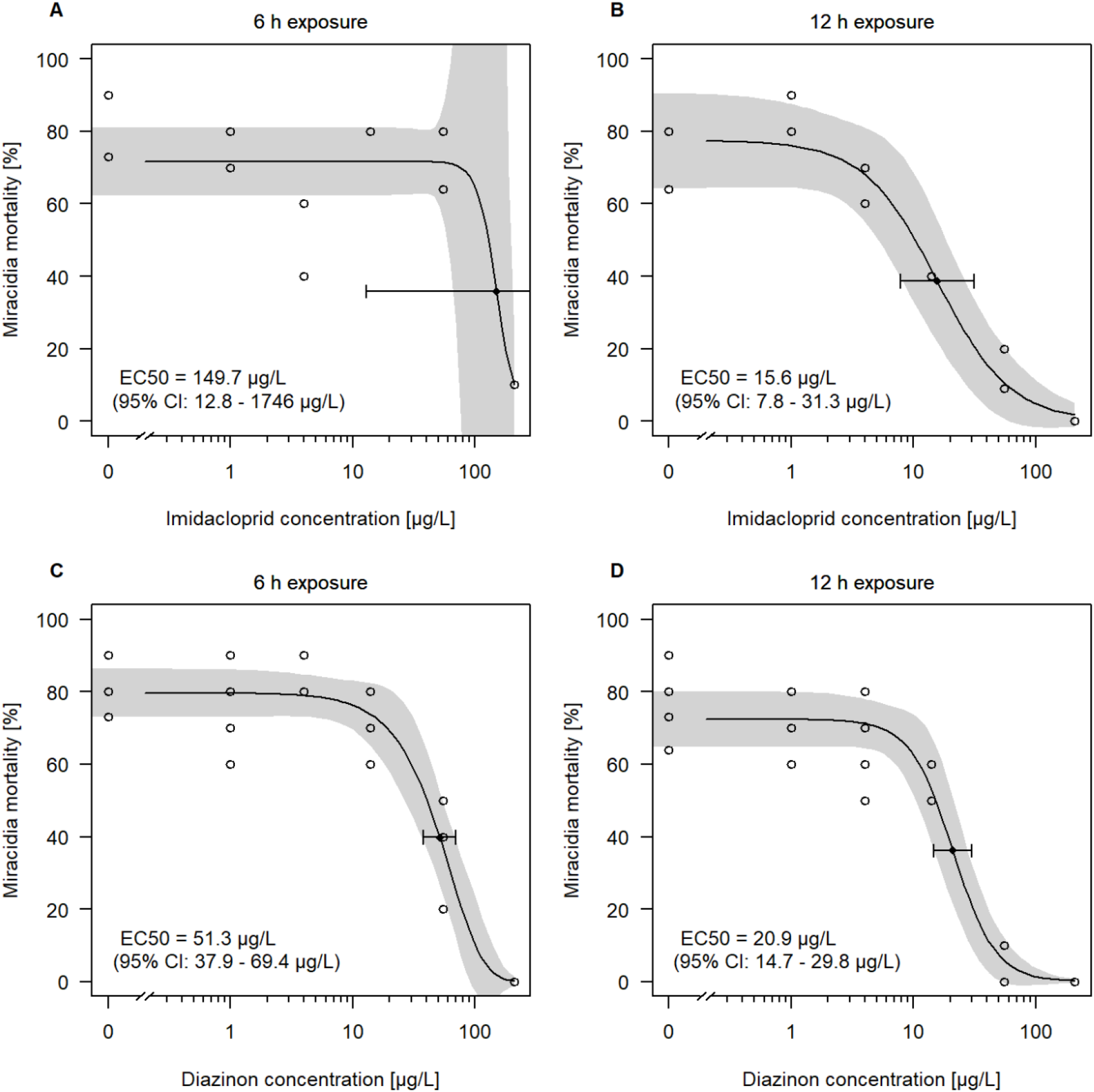
Immobilization of *S. mansoni* miracidia after exposure to (A, B) imidacloprid and (C, D) diazinon for 6 and 12 hours, respectively. Data points represent survival from different replicate tests, solid lines show the fitted concentration–response relationships, and the shaded areas correspond to the 95% confidence intervals. The EC50 (black dot) is shown together with the upper and lower limit of its associated 95 % confidence interval.

Cercariae were partly tested under ambient temperature conditions (ca. 25 °C) and under controlled conditions (18 °C). However, temperature had no significant effect on the concentration-immobilization relationship (*p* = 0.213, deviance = 1.75, d.f. = 1, residual d.f. = 126 for the comparison of quasi-binomial GLMs with and without the temperature:pesticide concentration interaction term during backward selection). Across both temperature regimes, diazinon showed a significantly steeper concentration-immobilization relationship than imidacloprid (concentration:pesticide interaction in the final model: *χ^2^* = 35.94; df = 1;*p* < 0.001). Therefore, tests with cercariae from both temperature regimes were merged in the following analysis using non-linear regression. Overall, cercariae showed greater tolerance than miracidia to imidacloprid, with an average median effective concentration (EC50) to imidacloprid of 403.0 (274.8 - 591.2) μg/L after 6 hours (Fig. 4A) and of 283.7 (206.0 – 390.7) μg/L after 12 hours (Fig. 4B). The tolerance of cercariae to diazinon was lower, with an EC50 of 24.8 (14.4 – 42.9) μg/L after 6 hours (Fig. 4C) and of 13.5 (7.7 – 23.5) μg/L after 12 h (Fig. 4D).

**Fig 4.**
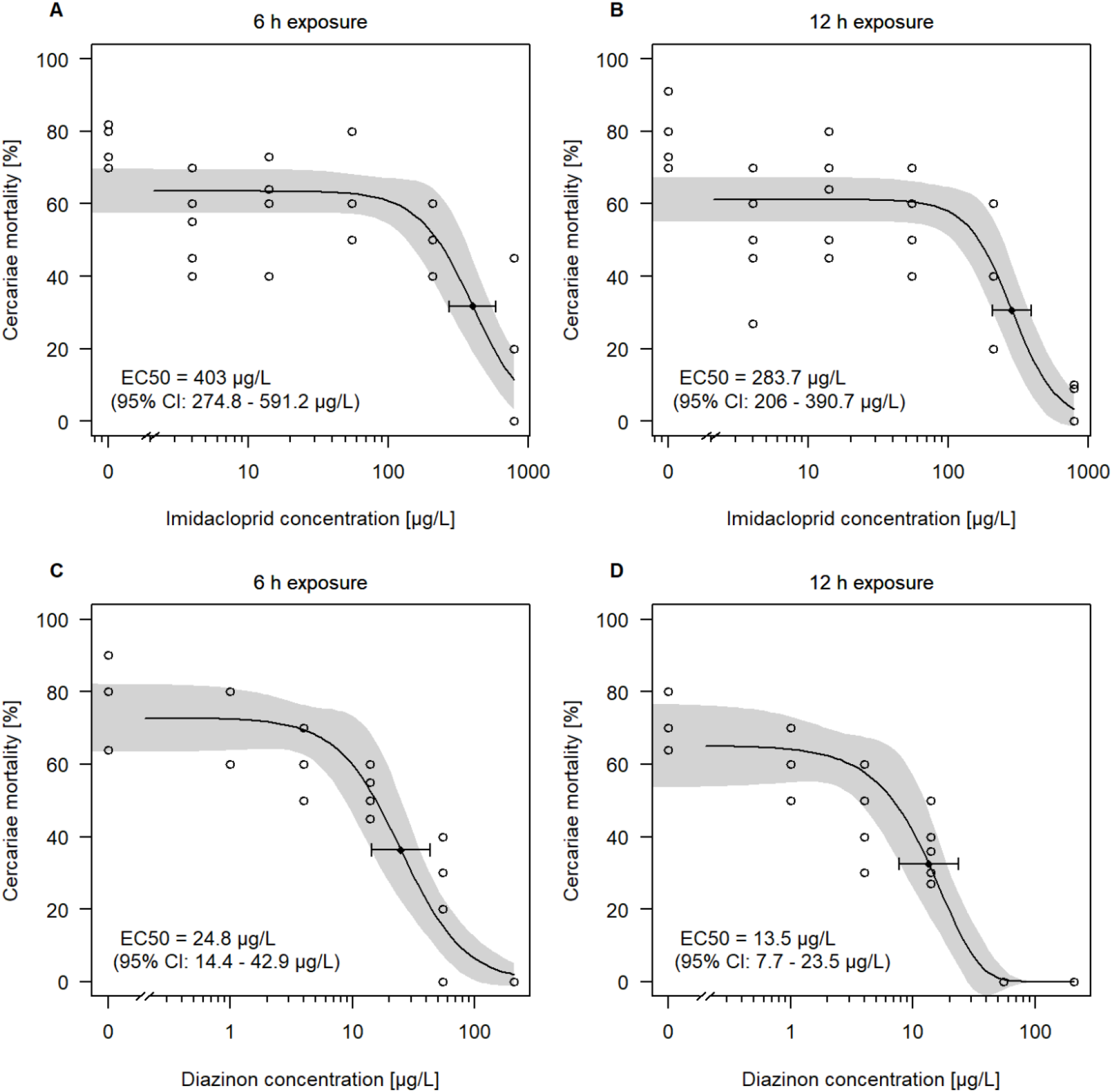
Immobilization of *S. mansoni* cercariae after exposure to (A, B) imidacloprid and (C, D) diazinon for 6 and 12 hours, respectively. Data points represent survival from different replicate tests, solid lines show the fitted concentration–response relationships, and the shaded areas correspond to the 95% confidence intervals. The EC50 (black dot) is shown together with the upper and lower limit of its associated 95 % confidence interval.

### Miracidia host-seeking assay

The effect of pesticide exposure on the infectivity of miracidia was identified based on the number of snails that were PCR positive for *Schistosoma* DNA (Tab. 2 in the supplementary materials). Miracidia were exposed to bottled water (control), low or high concentrations of imidacloprid or diazinon for up to two, four or six hours before they had access to a snail in bottled water. The low test concentrations,4.88 μg/L imidacloprid and 1.05 μg/L diazinon, equalled 3.3% and 2% of the 6h EC50 for miracidia, respectively. The high-test concentrations of 48.8 μg/L imidacloprid and 10.5 μg/L diazinon, equalled 33% and 20% of the 6h EC50 for miracidia, respectively. The low test concentrations caused less than 1%, and the high test concentrations caused less than 5% mortality of miracidia after exposure for six hours (< 6h EC5), according to the models for the acute toxicity tests (see Fig. 3). Pesticide identity (imidacloprid or diazinon) and exposure time (two, four and six hours) did not significantly affect the percentage of infected snails; they also did not significantly interact with the effect of pesticide treatment (control, low or high concentration) on the infectivity of miracidia. They were thus removed from our model. However, pesticide treatment significantly affected the infectivity of miracidia (*χ2* = 23.1, d.f. = 2,*p* < 0.001): While the low test concentrations showed no significant effect (*χ2* = 0.08, d.f. = 1, *p* = 0.774), the high test concentrations decreased the percentage of infected snails from 25.7% to 2.7% (*χ2* = 10.62, d.f. = 1, *p* = 0.002).

**Fig 5:**
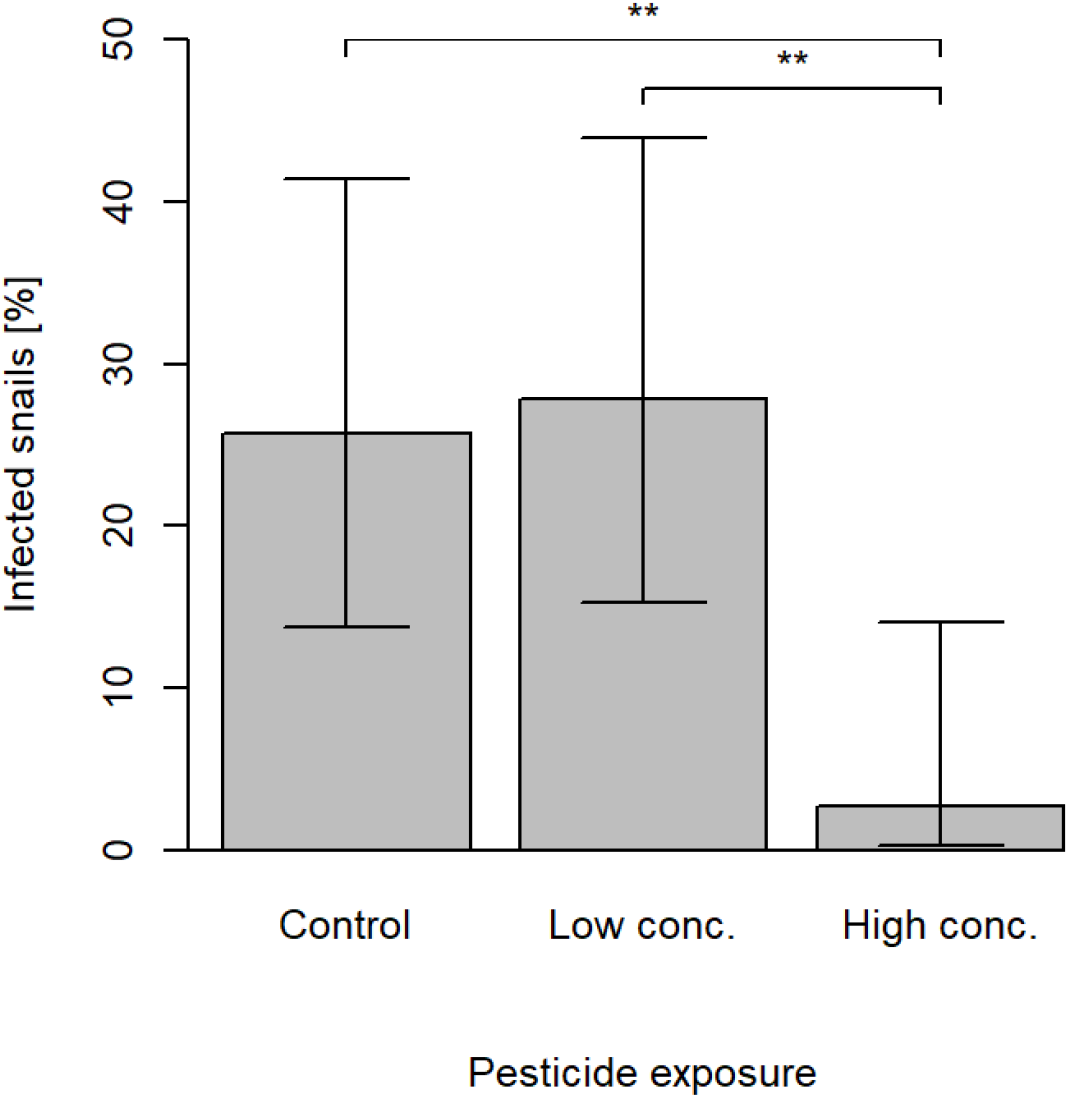
Percentage (mean and 95% confidence intervals) of infected snails after contact with five miracidia per snail for six hours. The miracidia had been previously exposed to either imidacloprid or diazinon for 2 – 6 h at low concentrations (imidacloprid: 4.88 μg/L; diazinon: 1.05 μg/L) that equalled 2 – 3.3 percent of the 6h EC50 for miracidia, and at high concentrations (imidacloprid: 48.8 μg/L; diazinon: 10.5 μg/L) that equalled 20 – 33 percent of the 6h EC50. Asterisks indicate significant changes in infection success as compared to the non-contaminated control (** *p* < 0.01).

### Sporocyst development assay

We tested the effect of weekly pesticide pulse exposure to the same nominal test concentrations as used in the miracidia host-seeking assay for 24 hours on the development of sporocysts within snails. Effects were identified based on the number of snails that had shed cercariae after 21 days (when the first snails started shedding) and after five weeks (Tab. 3 in the supplementary materials). No statistically significant effects could be observed. Overall, only 13 out of 118 snails (11%) shed cercariae after five weeks; the proportion of shedding snails did not differ between the controls (9%), low pesticide treatments (13%) and high treatments (12%; *χ^2^* = 0.25, d.f. = 2, *p* = 0.882 for the elimination of pesticide treatment as the last step in backward model selection).

The pesticide treatments did also not affect the survival of snails. Overall, 124 out of 280 snails (44 %) survived until the end of the test; survival did not significantly differ between the controls (48%) and low pesticide treatments (44%) or high treatments (40%; *χ* = 1.18, d.f. = 2, *p* = 0.555 for the elimination of pesticide treatment as the last step in backward model selection).

## Discussion

Our investigation showed that the maximum imidacloprid and diazinon concentrations observed in grab samples from natural water systems of the study area in Western Kenya [9] may not affect the aquatic life-stages of *S. mansoni* directly. The median effective concentrations that immobilized 50% of miracidia and of cercariae after constant exposure for 6 hours (6h EC50) were three to four orders of magnitude higher than the concentrations observed in the field. High, but sublethal concentrations of 48.8 μg/L imidacloprid and 10.5 μg/L diazinon reduced the infectivity of miracidia but not the development of sporocysts within host snails. However, these were 538 – 1,540 times higher than the concentrations observed in the aquatic environment [9]. Lower environmentally relevant concentrations of 4.8 μg/L imidacloprid and 1.05 μg/L diazinon showed no effects on the development of *S. mansoni*. The EC50s for the free-swimming *S. mansoni* life stages are within the same range as those for standard test organisms from the aquatic invertebrate community that are typically used in ecotoxicology: E.g., the 6h EC50s of imidacloprid for miracidia and cercariae were 2.7 and 7.3 times as high as the acute 96h EC50 for *Chironomus riparius* (55 μg/L) [28]. The 6h EC50s of diazinon for miracidia and cercariae were 51 and 25 times as high as the 48h EC50 for *Daphnia magna* (1 μg/L) [28]. The EC50 values are not directly comparable as the exposure times in the tests differ, such that a simple ranking may overestimate the relative tolerance of *Schistosoma*.

Nevertheless, our results suggest that *S. mansoni* is likely to benefit indirectly from pesticide exposure because it can survive together with its host snails in waters where competitors and predators of the snails cannot. This conclusion is based on three considerations: First, in contrast to other invertebrates, *S. mansoni* may be rarely exposed to harmful concentrations because its free-swimming life stages are only present in the water column for short periods of time. Results from the sporocysts development assay indicate that after successful infection, *S. mansoni* is well protected from pesticide exposure within the tissue of its host snail. In the field, accumulation of pesticides and thus exposure of sporocysts within the snails might be higher than in the experiment due to additional exposure pathways such as contaminated snail food and because snails may accumulate pesticides already before infection. Nevertheless, in the form of sporocysts, *S. mansoni* can escape high short-term pesticide exposure peaks to the water that follow run-off events and drive the overall risk of pesticides to invertebrates in agricultural streams [14]. Second, the standard test organisms in ecotoxicology do not always represent the most sensitive macroinvertebrate species that live in freshwaters. In acute tests, some potential competitor or predator species of host snails such as ephemeropteran species appeared about 50% less tolerant to the tested insecticides [6]. Moreover, our tests covered all relevant aquatic life stages of *S. mansoni*, whereas acute toxicity tests are limited to a single life stage that may not be the most sensitive one. Particularly for insects it has been shown that larvae can survive concentrations in acute tests that result in considerably higher delayed mortality during moulting [29, 30]. Therefore, particularly insects can be more sensitive than they appear from the available acute toxicity tests, while this seems unlikely for *S. mansoni*.

Third, the very high pesticide tolerance of host snails renders indirect effects on sporocyst development due to disturbed nutrient supply within snails unlikely also under field conditions. The observed 6h EC50s of imidacloprid for the miracidia and cercariae of *S. mansoni* for imidacloprid fall more than 2,400 times lower than the 24h EC50 for the highly tolerant host snail *Bulinus pfeifferi* (> 1 g/L) [6]. The 6h EC50s of diazinon for miracidia and cercariae fall more than 390 times lower than the 24h EC50 for *B. pfeifferi* (~ 20 mg/L) [6]. Given this high tolerance of *B. pfeifferi*, it is not surprising that we also did not observe an indirect positive effect of pesticides on the infectivity of miracidia that could arise from adverse effects on the host snail immune system.

It should be noted that in the field, macroinvertebrates can respond negatively to pesticides at considerably lower concentrations than in the laboratory [6,31]. This can be related to factors such as additional stressors [32,33] and mixture toxicity [34]. Nevertheless, the observed no-effect concentrations (NOEC) of 4.88 μg/L imidacloprid and 1.05 μg/L diazinon for the overall performance of *S. mansoni* throughout its aquatic life stages were still around 60 times higher than the maximum environmental concentrations observed in western Kenya [9]. For comparison, NOECs in the aquatic environmental risk assessment of pesticides are often divided by an empirically obtained factor of 10 – 100 in order to account for variability in sensitivity across both species and environmental conditions [34]. However, with information on inter-species variability available, this assessment factor can be lowered e.g. in Europe to 3 – 6 to account only for variability across environmental conditions [35]. Though the protectiveness of assessment factors has been questioned [36] the lowered factors are an order of magnitude lower than the ratio of our observed NOEC and the environmental concentrations in western Kenya. Therefore, we expect no considerable effects of pesticide exposure on the performance of *S. mansoni* to occur also under natural conditions.

We used field-collected snails that were reared for several weeks prior to the experiments to ensure they were free of schistosome infections. Comparably high mortality during breeding and in the controls indicated suboptimal breeding and test conditions, and limited the number of available replicates. This should be considered when comparing the obtained EC50 values with those from standard tests under optimum conditions. Nevertheless, suboptimum conditions represent additional stress that is likely to increase rather than decrease effects of pesticides (see discussion above). Therefore, we consider the risk of having missed relevant effects due to suboptimum test conditions low.

Taken together, our results suggest that exposure of surface waters to agricultural pesticides may indirectly increase the risk of schistosomiasis transmission by sparing the pathogen *S. mansoni*, while the natural control of its host snail *B. pfeifferi* is affected [6]. Thus, pesticide mitigation measures should be taken in at-risk areas to prevent further exacerbation of the disease.

## Acknowledgement

This work was supported by the DFG (Deutsche Forschungsgemeinschaft, https://www.dfg.de/en/), grant number LI 1708/4-1, HO 3330/12-1 and the German Helmholtz long-range strategic research funding. We also gratefully acknowledge the financial support for this research by icipe’s core donors, the Swedish International Development Cooperation Agency (Sida, https://www.sida.se/en); the Swiss Agency for Development and Cooperation (SDC, https://www.eda.admin.ch/sdc); Federal Democratic Republic of Ethiopia; and the Kenyan Government.

## Supplementary Information

**Fig S1:**
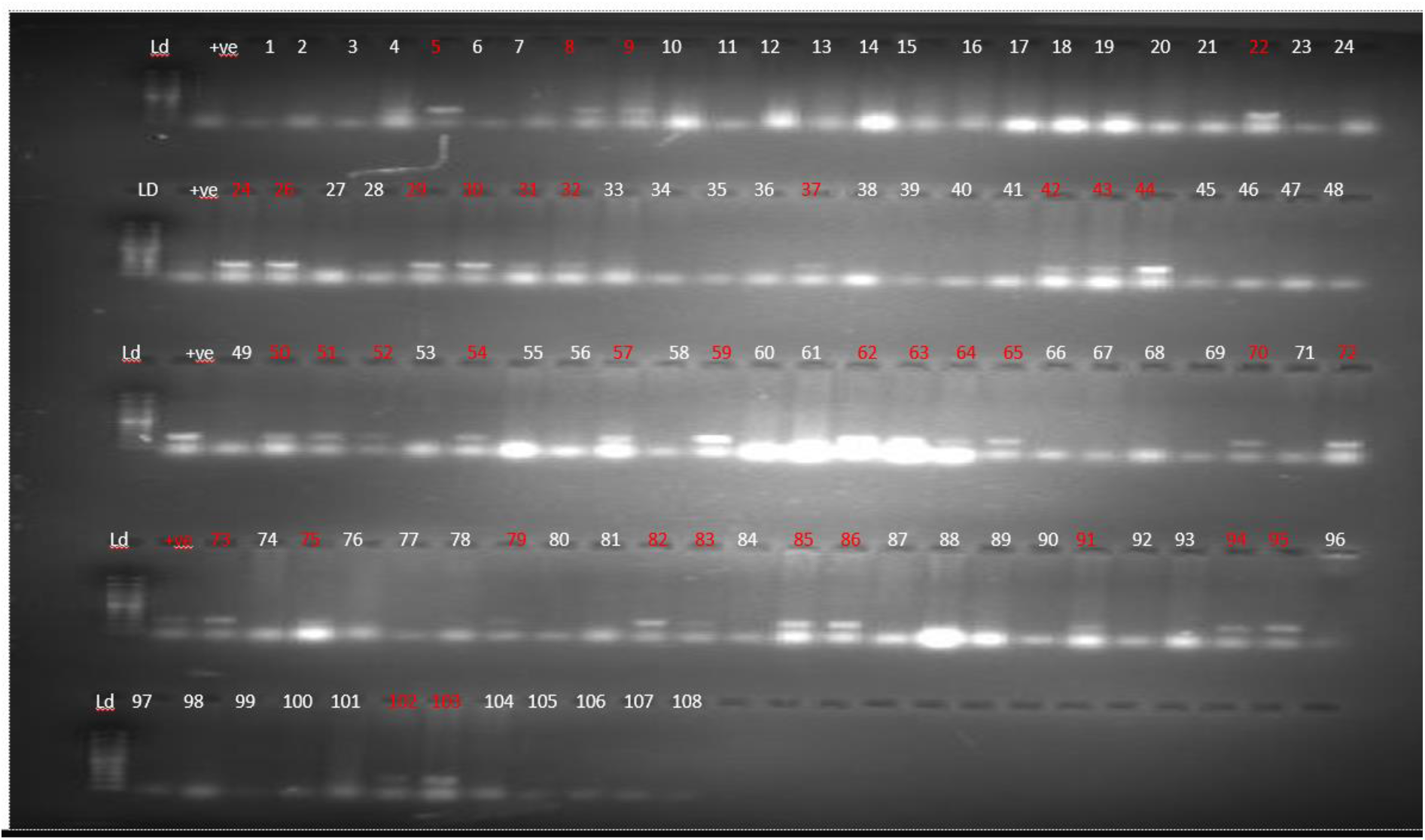
Gel image of the PCR results of the miracidia host seeking assay for imidacloprid. The image shows the results for controls (well 1-12= controls in substrate for 2 hours, 13-24= controls in substrate for 4 hours and 24-36= controls in substrate for 6 hours) versus those exposed to imidacloprid at 2% the average EC50 for miracidia at 6 hrs (37-72, arranged for time as controls) and those exposed to imidacloprid at 40% the average EC50 for miracidia at 6 hrs (73-108). Each row starts with a DNA ladder and a positive control (250 bps) except the bottom row which does not have the positive control.

**Table S2:** Results of the miracidia host seeking assay in raw format.

| Pesticide | Treatment | Concentration | Infected | Not infected | Percentage infected |
| --- | --- | --- | --- | --- | --- |
| Control | Control | 0 | 3 | 9 | 25.86 |
| Control | Control | 0 | 2 | 10 |  |
| Control | Control | 0 | 5 | 8 |  |
| Control | Control | 0 | 5 | 7 |  |
| Control | Control | 0 | 4 | 8 |  |
| Control | Control | 0 | 0 | 13 |  |
| Imidacloprid | Low | 4.88 | 4 | 8 | 44.44 |
| Imidacloprid | Low | 4.88 | 6 | 6 |  |
| Imidacloprid | Low | 4.88 | 6 | 6 |  |
| Imidacloprid | High | 48.8 | 0 | 12 | 2.78 |
| Imidacloprid | High | 48.8 | 0 | 12 |  |
| Imidacloprid | High | 48.8 | 1 | 11 |  |
| Diazinon | Low | 1.05 | 1 | 11 | 11.11 |
| Diazinon | Low | 1.05 | 0 | 12 |  |
| Diazinon | Low | 1.05 | 3 | 9 |  |
| Diazinon | High | 10.5 | 0 | 12 | 2.78 |
| Diazinon | High | 10.5 | 0 | 12 |  |
| Diazinon | High | 10.5 | 1 | 11 |  |

**Table S3:** Results of the sporocyst development assay in raw format.

| Pesticide | Treatment | Concentration | Shedding | Not shedding | Percentage shedding | Percentage shedding on week 1 |
| --- | --- | --- | --- | --- | --- | --- |
| Control | Control | 0 | 1 | 7 | 21.03 | 8.46 |
| Control | Control | 0 | 1 | 5 |  |  |
| Control | Control | 0 | 1 | 5 |  |  |
| Control | Control | 0 | 1 | 6 |  |  |
| Control | Control | 0 | 2 | 5 |  |  |
| Control | Control | 0 | 3 | 5 |  |  |
| Imidacloprid | Low | 4.88 | 3 | 5 | 23.61 | 8.33 |
| Imidacloprid | Low | 4.88 | 0 | 8 |  |  |
| Imidacloprid | Low | 4.88 | 2 | 4 |  |  |
| Imidacloprid | High | 48.8 | 2 | 2 | 33.33 | 11.11 |
| Imidacloprid | High | 48.8 | 0 | 3 |  |  |
| Imidacloprid | High | 48.8 | 3 | 3 |  |  |
| Diazinon | Low | 1.05 | 0 | 3 | 26.67 | 12.5 |
| Diazinon | Low | 1.05 | 4 | 1 |  |  |
| Diazinon | Low | 1.05 | 0 | 4 |  |  |
| Diazinon | High | 10.5 | 2 | 2 | 27.78 | 11.11 |
| Diazinon | High | 10.5 | 2 | 4 |  |  |
| Diazinon | High | 10.5 | 0 | 7 |  |  |

